# Glucocorticoid receptors mediate reprogramming of astrocytes in depression

**DOI:** 10.1101/2025.04.04.646549

**Authors:** Sedef Dalbeyler, Aleksandra Herud, Bartosz Zglinicki, Patrycja Ziuzia, Marcin Piechota, Dzesika Hoinkis, Laura Bergauer, Carmen Menacho Pando, Zbigniew Soltys, Paweł Hanus, Patrick Groves, Luanna Dixon, Florian Ganglberger, Dolores Del Prete, Valentyna Dubovyk, Marlene Aßfalg, Coralie Violet, Nathan Lawless, Szymon M. Kielbasa, Hannes Sigrist, Christopher Pryce, Gustavo Turecki, Mathias V. Schmidt, Vladimir Benes, Thomas Kuner, Michal Korostynski, Bastian Hengerer, Michal Slezak

**Author notes:** **Correspondence** Dr Michał Ślęzak, Biology of Astrocytes Research Group, Lukasiewicz Research Network – PORT, Polish Center for Technology Development, Wroclaw, Poland., Stabłowicka 147, 54-066 Wroclaw, Poland. These authors contributed equally (Sedef Dalbeyler, Aleksandra Herud, Bartosz Zglinicki, Patrycja Ziuzia).

## Abstract

Psychiatric disorders are among the most pressing problems of the modern society, with various forms of depression affecting more than 300 millions of people worldwide. Dysfunction of glial cells has consistently been reported in major depressive disorder (MDD); however, no comprehensive resource detailing glial dysfunction is available. To provide insight into neurobiological mechanisms behind severe psychiatric symptoms, we performed transcriptional analysis of post-mortem samples from a subpopulation of suicide completers with previously reported glial abnormalities. We focused on BA25, a subregion of the prefrontal cortex prioritized for targeted medical interventions, due to its metabolic aberrations in disease. We found that a significant portion of genes deregulated in MDD is enriched in glia, with astrocyte-specific genes representing the highest fraction. Then we employed a novel protocol for enriching astrocytic nuclei to provide a detailed molecular signature of astrocytes in MDD. The analysis of the gene set revealed the glucocorticoid receptor (GR) as a key regulatory transcription factor. We found that astrocyte-specific elimination of the GR in mice largely prevented transcriptional, metabolic and behavioral changes elicited by chronic stress. We also demonstrated that regional manipulation of glutamate turnover in astrocytes suffices to elicit discrete traits of depressive-like behavior. Our data points to astrocytes as a key cellular site of convergence of multiple traits of depression and provide a resource for exploring novel targets for glia-focused therapeutic approaches.

## INTRODUCTION

Mental disorders are heterogenous with respect to their etiology, manifestations and response to treatment, which makes them a major cause of global disability^1,2^. At the most extreme scenario, mental health problems may lead to suicide, a leading cause of death among young people^3^. Modern research frameworks aim to resolve the biological background of individual symptoms of mental disorders for improved patient stratification and precise therapies^4,5^. This effort is being accomplished through the integration of data on dysfunction of neural circuits underlying behavioral symptoms with biochemical markers and risk factors^6,7^. The most important risk factor of developing major depression is stress, experienced as an early childhood trauma or chronically^8,9^. At the molecular level, the stress response is coordinated by the hypothalamus-pituitary-adrenal (HPA) axis, with the most important effectors being glucocorticoids (GCs)^10^. It has been shown that genetic variation in gene networks regulated by glucocorticoid receptor (GR) in humans confers the risk for psychiatric diseases and predicts the outcome of antidepressant treatment^11,12^. Physiologically, GCs act as a molecular effector of the HPA axis activity, which shapes systemic metabolism through the regulation of circadian transcriptional program in highly cell type-specific fashion^13^. While mechanisms linking GR signaling route exploited for adaptive learning and memory are described^14^, dysfunctions of neural circuits at the level of cell types and molecular pathways associated with psychiatric conditions has only started to be discovered.

Astrocytes control brain energy metabolism and multiple aspects of synaptic transmission^15,16^. Altered expression of astrocyte-specific genes crucial for these processes was repetitively reported in transcriptional surveys of post-mortem samples of various brain regions collected from depression and suicide cases^17-20^, particularly in male subjects^21^. Despite these premises, astrocytes have gained very limited attention in data-driven therapeutic pipelines. This surprising gap largely stems from the lack of knowledge on a disturbance of astrocytes in depression.

In depression, several brain regions display corrupted metabolic activity, including subgenual prefrontal cortex^22,23^. Notably, reversal of these abnormalities correlates with the release of depressive symptoms upon targeted intervention^24,25^. One of the relevant regions was Brodmann area 25 (BA25), transcriptionally altered in depression and suicide^26,27^. Due to its role in the stress response^28^, emotion processing^29^, and the regulation of systemic metabolism^30^, we aimed for a detailed characterization of astrocyte deficiency from this brain region. Driven by the outcome of the initial analyses, we performed a systematic investigation of the impact of the main risk factor of depression, stress, and its main systemic mediators, GCs, on the molecular profile and the function of astrocytes.

## RESULTS

### Downregulation of glia-specific genes in BA25 of ‘low expressors’

To explore glial deficiency in severe psychiatric conditions, we performed the transcriptional profiling of BA25 from a unique and well described resource. We used a collection containing brain samples of suicide completers, who were diagnosed with major depressive disorder (MDD). A previous study showed, that in approximately one third of 74 MDD samples, the expression of some astrocyte-specific genes was decreased in several brain regions. Based on these changes, respective subjects were classified as ‘low expressors’^18^. We reasoned that since this subpopulation represents the extreme example of astrocytes’ transcriptional impairment, thorough inspection of these samples shall allow a detailed insight into aberrant molecular pathways specific for astrocytes.

To examine the insight into the molecular pathology of these specimens, we generated single nuclei homogenates of BA25 samples dissected either from ‘low expressors’^1^ (MDD, n = 15) or controls (CTRL, n = 12), the latter group encompassing cases of accidental or natural death (Suppl. File 1, Table S1). We fluorescently labeled nuclei with a nuclear dye Hoechst and sorted them for subsequent transcriptional profiling (Fig. 1A, ‘Hoechst+’, Suppl. Methods). We found no differences in the number of labeled nuclei nor the quantity and quality of the extracted genetic material (Fig. S1).

**Fig. 1.**
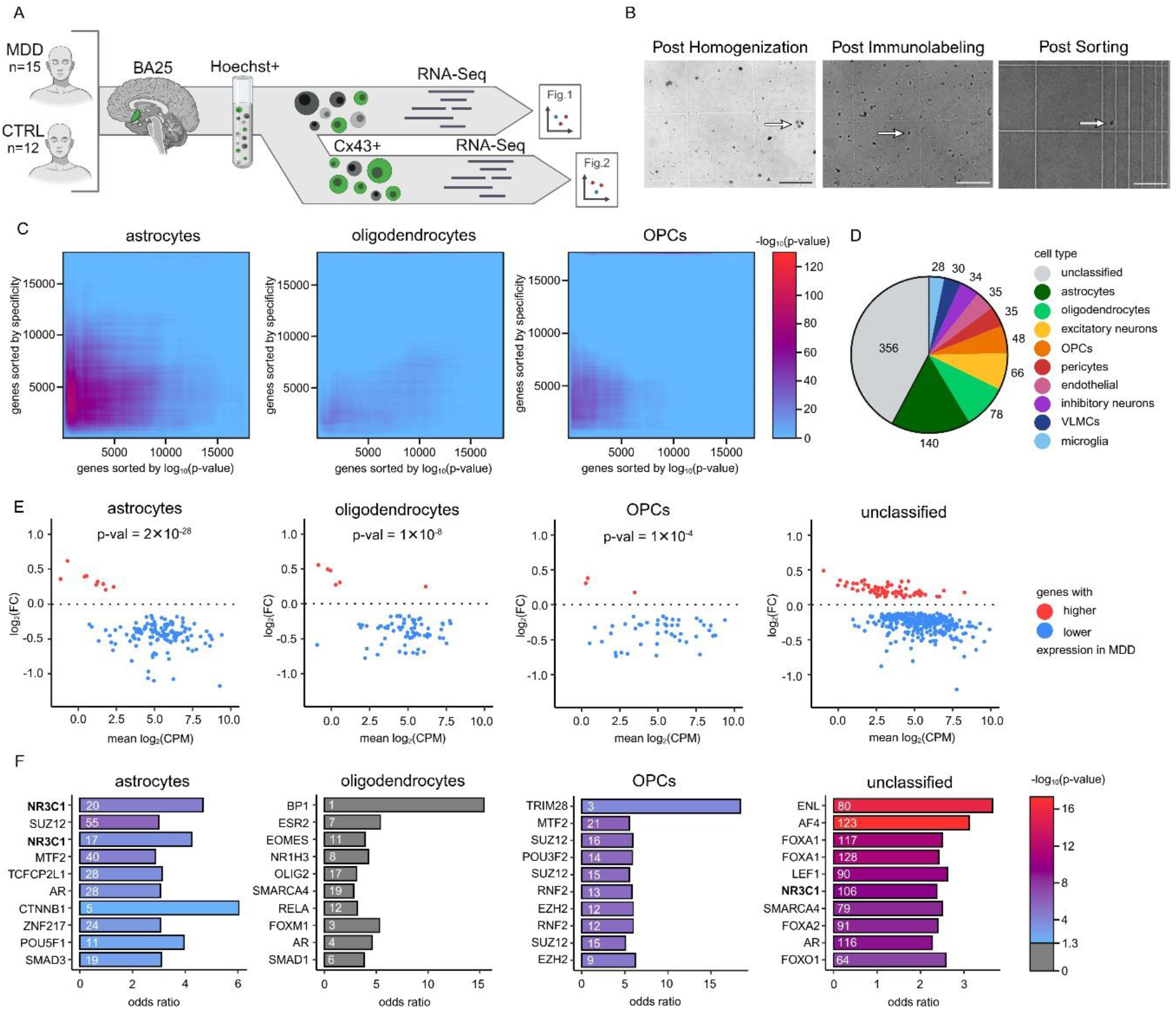
Glucocorticoid receptor is a key regulatory factor of astrocytes’ reprogramming in MDD. **A**. Graphical representation of the experimental workflow. BA25 samples were dissected from a subset of brains of suicide completers diagnosed with MDD and control cases. Hoechst+ and Cx43+ nuclei were isolated from the tissue and RNA sequencing was performed. **B**. Microphotographs of nuclei after tissue homogenization, immunolabeling and sorting. Scale bar, 125 μm. **C**. Rank-rank hypergeometric overlap constructed with the list of genes ranked by cell type specificity and detected genes log_10_(p-value). Color scale represents -log_10_(p-value) of the overlap. **D**. Distribution of Hoechst DEGs based on their assignment to specific cell types. **E**. Distribution of log_2_(FC) and mean log_2_(CPM) of DEGs showing significant overlap with cell type-specific gene lists and DEGs not assigned to any cell type (‘unclassified’). **F**. Top 10 results of transcription factors over-representation analysis performed for cell type-specific DEGs and ‘unclassified’ DEGs, sorted by a combined score. Color scale represents -log_10_(p-value) of the overlap between TF-specific datasets and cell-type specific DEGs. Values on the bars indicate a number of overlapping genes.

To examine the contribution of cell types to MDD-related variance, we ranked all detected protein-coding genes by the p-value of differential expression between MDD and CTRL samples and using Rank Rank Hypergeometric Overlap (RRHO), we examined the list against lists of the same genes ranked by the parameter reflecting their enrichment in each of the major brain cell types, normalized with BrainTrawler, based on the single cell sequencing data from the human cortex^2^ (Suppl. Methods). This analysis revealed significant agreement for astrocytes, oligodendrocytes and oligodendrocyte progenitor cells (OPCs) (Fig. 1B).

Subsequent analysis revealed 843 differentially expressed genes (DEGs) between MDD and CTRL at the nominal p-value < 0.05, out of which 578 showed decreased expression in MDD samples (Suppl. File 1, Table S2). This transcriptional signature was again examined for outstanding contribution of major brain cell types and upstream regulators.

To test for the association of observed DEGs with specific cell types, we examined the DEGs list using ENRICHR. We found striking enrichment of genes specific for glial cells: astrocytes (e.g. ‘Human Astro L1 FGFR3 SERPINI2 up’, adj. p = 5.3 x 10^-46^) and oligodendroglial lineage (e.g. ‘Human Oligo L3-6 OPALIN ENPP6 up’, adj. p = 3.2 x 10^-28^; ‘Human OPC L1-6 PDGFRA COL20A1 up’, adj. p = 6.5 x 10^-13^) (Suppl. File 1, Table S3). Direct testing of Hoechst+ DEGs against normalized signatures of nine major brain cell types, confirmed significant overlap with ‘Astrocytes’ (adj. p = 2.4 x 10^-28^), ‘Oligodendrocytes’ (adj. p = 1.5 x 10^-8^) and ‘OPCs’ (adj. p = 1.1 x 10^-4^), while a large fraction of DEGs was not assigned to any specific cell type (Fig. 1B, Suppl. File 1, Table S4-S13). Vast majority of glia-specific DEGs were downregulated in MDD samples (Fig. 1C, Suppl. Fig. 2). This data demonstrated that transcriptional reprogramming in BA25 of ‘low expressors’ occurs primarily in glial cells, with the most prominent effects observed in astrocytes. In fact, 11 most downregulated DEGs in ‘low expressors’ were astrocyte-specific (Suppl. File 1, Table S2).

### Glucocorticoid receptor is an upstream regulatory factor of astrocyte-specific DEGs

Next, we explored upstream regulators of cell type-specific DEGs. To this end, we performed over-representation analysis (ORA) of transcription factors (TFs) in sets of cell-type specific and unclassified DEGs, using ChEA 2022 database^3^ (Suppl. File 1, Table S15-S18). For astrocyte-specific DEGs, the analysis revealed significant overlap with 87 genesets, with *NR3C1*, encoding glucocorticoid receptor (GR), showing highest combined score (Fig. 1D, Suppl. File 1, Table S15). Notably, *NR3C1* was also overrepresented in ‘unclassified’ pool of DEGs, i.e. expressed in multiple cells types, including glia, as were receptors of other hormones (AR, ESR1, ESR2). For oligodendrocyte-specific DEGs, none of the TFs passed the statistical criteria for the overlap. The TFs enriched in the ‘OPCs’ dataset shared with ‘astrocytes’ included members of the polycomb group protein regulating cell differentiation and maturation, e.g. *SUZ12*, and *MTF2*. This data highlighted the importance of hormonal signaling for the neuropathology of severe depression and suggested a key role of the GR for astrocyte reprograming in ‘low expressors’.

### Deregulation of metabolic and synaptic pathways in astrocytes of ‘low expressors’

To gain a detailed insight into alterations of molecular pathways in astrocytes from ‘low expressors’, we established a protocol enabling the enrichment of astrocytic nuclei from fresh-frozen samples of human brain (Fig. 1A; Suppl. Methods). Positive selection was accomplished with the antibody recognizing a protein specific for astrocytes in the adult brain, CX43, encoded by the gene *GJA1*. The protein is enriched in the plasma membrane and the endoplasmatic reticulum, the latter known to attach to nuclei in tissue homogenates^6^. We applied the enrichment protocol to Hoechst+ nuclei suspension of the original set of post-mortem BA25 samples and subjected resulting nuclei fraction (‘Cx43+’) to RNA-Seq.

The gene set enrichment analysis (GSEA) of gene sets derived from the GO Biological Process ontology examined all genes detected in astrocyte nuclei RNA-Seq and revealed the overrepresentation of crucial pathways engaged in glutamate turnover, fatty acid metabolism, amino acid transport, development, and synapse organization (Fig. 2A, Suppl. File 2, Table S5).

**Fig. 2.**
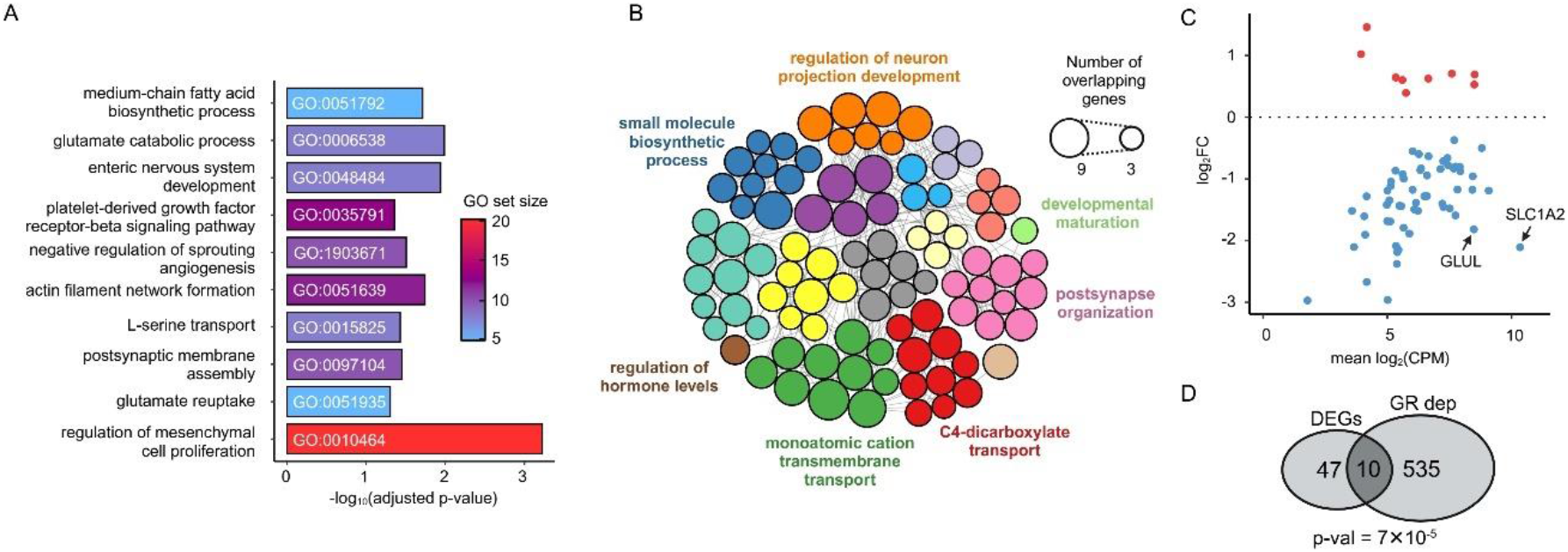
GR-dependent network controls reprogramming of astrocytes in BA25 ‘low expressors’. **A**. Top 10 results of the Gene Set Enrichment Analysis sorted by rank parameter. Color scale represents GO set size. **B**. Over-representation analysis of protein-coding DEGs, visualized as a network consisting of GO BP terms grouped into clusters named with the most significant GO term in the cluster. Size of the circles represents number of genes overlapping between genes in the GO term and Cx43+ DEGs. **C**. Distribution of log_2_(FC) and mean log_2_(CPM) of Cx43+ DEGs (MDD vs CTRL). **D**. The overlap of Cx43+ DEGs with astrocyte-specific, GR-dependent geneset.

The subsequent analysis revealed 67 protein-coding DEGs that passed the stringent statistical criteria (FDR < 0.1, resulting in p < 0.00041) in the ‘Cx43+’ nuclei population (Suppl. File 2, Table S1). In this set, 58 DEGs displayed lower expression in MDD samples. None of the genes showing higher expression in MDD samples displayed high specificity for astrocytes (Suppl. Methods). The ORA performed on the ‘Cx43+’ DEGs confirmed observations from the GSEA, including the regulation of synaptic transmission (e.g. ‘postsynapse organization’), metabolism (e.g. ‘small molecule biosynthetic process’) and transmembrane transporters (e.g. ‘C4-dicarboxylate transport’) (Fig. 2B, Suppl. File 2, Table S3-S4). Notably, detailed inspection of the list of ‘Cx43+’ DEGs, revealed that key regulators of glutamate uptake and turnover were among most downregulated genes: *SLC1A2*, encoding glia-specific glutamate transporter GLT1, responsible for the take up of excessive extracellular glutamate and *GLUL* encoding glutamine synthetase, an enzyme converting glutamate to glutamine, known to be a main route of glutamate turnover between astrocytes and neurons^31^ (Fig. 2C).

To independently test the GR regulation of astrocyte-specific DEGs, we compared the ‘Cx43+’ DEGs with a published resource of GR-dependent regulatory network in astrocytes. Since GR signaling is highly cell type-specific and no data existed for GR-dependent genes in human astrocytes, we used the comprehensive list of genes regulated by the exposure to GR agonist in primary mouse astrocytes^7^. We found significant overlap between the ‘Cx43+’ DEGs and ‘GR-dependent regulatory network in astrocytes’ (Fig. 2D). The shared pool included *SLC1A2* and *GLUL*, as well as several genes known to mediate crucial metabolic functions of astrocytes (e.g. *ALDOC, NT5E, DIO2*) (Suppl. File 2, Table S2).

In sum, these analyses indicated a breakdown of fundamental pathways operating in astrocytes in the BA25 of ‘low expressors’ and its regulation by the GR.

### GR controls CSDS-induced transcriptional reprograming of astrocytes in mice

Next, we tested the role of the GR in mediating reprograming of astrocytes in an ethologically relevant rodent model. We employed the chronic social defeat stress (CSDS) paradigm, broadly used in preclinical studies to induce sustainable physiological and behavioral symptoms of depressive-like phenotype in mice^32^ (Fig. 3A, Fig. 5). For selective elimination of the GR from astrocytes, we bred a transgenic line carrying an exon 3 of the *Nr3c1* gene flanked by loxP sites with Aldhl1-CreER^T2^ driver line. Bigenic mice (GR^astroKO^) and their monogenic littermates (GR^lox/lox^) were treated with TAM, which resulted in halving the fraction of astrocytes with detectable expression of the GR protein in GR^astroKO^ animals, keeping neuronal GR expression unaffected (Fig. 3B, Suppl. Fig. 3). Animals were assigned to two groups: the control group (CTRL), where mice were housed in pairs, and the CSDS group, were social defeat was accomplished for 15 days (Fig. 3A). Upon completion of the CSDS, all mice were sacrificed and astrocytes were isolated by magnetic cell sorting (MACS), and subjected to RNA-Seq (Suppl. Methods). For the mechanistic follow-up of the human data, we investigated astrocytes from the prefrontal cortex (PFC), which includes areas analogous to human BA25, and the hippocampus (HIP), as both are key nodes in the neurocircuitry mediating the effects of chronic stress and relevant to depression-like phenotypes.

**Fig. 3.**
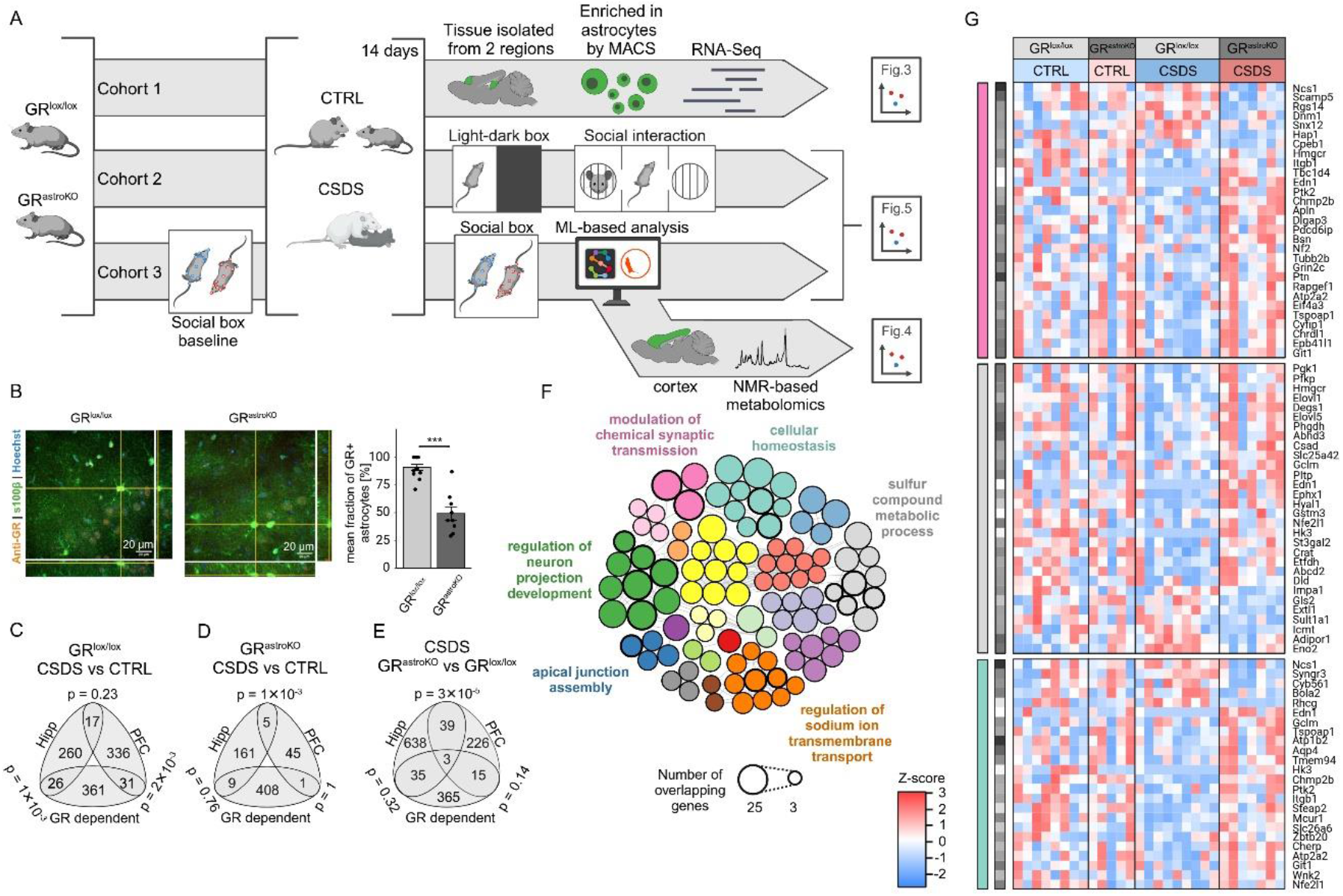
GR-dependent signaling controls molecular reprogramming of astrocytes in CSDS. **A**. Graphical scheme of the workflow. **B**. Example micrographs from immunohistochemical analysis of the GR expression in the PFC of GR^lox/lox^ and GR^astroKO^ mice and barplots summarizing the fraction of cells double labelled against GR and s100β. p***<0.001, t-test. **C-E**. Venn diagrams picturing overlaps between DEGs from PFC or HIP, and GR-dependent genesets. **F**. Over-representation analysis of significant protein-coding genes of GR^astroKO^ /CSDS vs GR^lox/lox^/CSDS comparison in the PFC, visualized as a network consisting of GO BP terms grouped into clusters named with the most significant GO term in the cluster. Circles in bold represent GO terms shared between human MDD vs CTRL in Cx43+ DEGs and mouse GR^astroKO^/CSDS vs GR^lox/lox^/CSDS comparison. Size of the circles represent number of genes overlapping between genes in the GO term and DEGs of GR^astroKO^/CSDS vs GR^lox/lox^/CSDS. **G**. Heatmaps showing expression levels of genes belonging to selected GO BP clusters from F, and significant in GR^astroKO^/CSDS vs GR^lox/lox^/CSDS in the PFC. Black and white vertical bar represents mean transcript abundance.

The statistical analysis revealed major impact of the CSDS on astrocytes isolated from GR^lox/lox^ mice, with 488 DEGs in the PFC (268 downregulated) and 413 DEGs in HIP (186 downregulated) (p < 0.05, GR^lox/lox^/CSDS vs GR^lox/lox^/CTRL). The minimal overlap between the two examined regions (Fig. 3C) indicated high regional specificity of GR-regulated transcription in astrocytes. The comparison with the list of astrocyte-specific, GR-dependent genes revealed significant overlap with astrocyte-specific DEGs in both, the PFC (p = 2 × 10^-3^) and HIP (p = 1 × 10^-3^) (Fig. 3C). These transcriptional effects of CSDS were dramatically reduced by astrocyte-specific elimination of the GR. The number of region-specific DEGs was much lower in GR^astroKO^ mice (PFC: 64 DEGs, HIP: 240 DEGs), as was the overlap between the two regions. Additionally, no statistically significant overlap with GR-dependent geneset was detected in GR^astroKO^ astrocytes from any region (Fig. 3D).

Finally, we examined the CSDS-induced transcriptional response differentiating astrocytes from GR^lox/lox^ and GR^astroKO^ in both brain regions. In this comparison, we found 345 DEGs in the PFC and 941 DEGs in HIP (p < 0.05, GR^astroKO^/CSDS vs GR^lox/lox^/CSDS) (Fig. 3E). This data confirmed that GR signaling regulated the impact of CSDS on the molecular profile of astrocytes (Suppl. File 3, Table S1-S6).

### Astrocyte-specific dysfunction in PFC shared between human MDD and mouse CSDS

Next, we examined which CSDS-induced pathways relied on GR signaling. Due to the complementary data from human samples, we focused on PFC astrocytes. We analyzed differential GOs regulated by CSDS in GR^astroKO^ and GR^lox/lox^ animals (Fig. 3F). The ORA performed on the set of DEGs revealed 118 GO differential pathways (Suppl. File 3, Table S8-S10). Sixteen of those matched GOs differentiating CTRL and MDD human Cx43+ nuclei from BA25 (compare Fig. 3F and 2C, bold circles).

Detailed inspection of differential GOs revealed that they belong to three main groups: pathways engaged in synaptic communication (example: Fig. 3G, pink cluster), metabolism (example: Fig. 3G, grey cluster) and cellular homeostasis (example: Fig. 3G, teal cluster). Further examination of genes contributing to the overlap revealed a handful of hits shared between the human ‘Cx43+’ DEGs and mouse PFC DEGs, including: *Chrdl1*, known to play a role in synapse formation, *Grin2c*, encoding a subunit of the NMDA receptor enriched in astrocytes, *Atp1b2*, encoding a Na+-K+ transporting ATPase and *Ptn*, encoding pleiotrophin, a secreted heparin-binding growth factor. These genes are hence considered translational markers of depressive state of the brain.

### Disturbed glutamate turnover in astrocytes controls circuit-specific behavior

Next, we examined the metabolic effects of CSDS in the brain. To this end, we homogenized cortices obtained from a new cohort of CTRL and CSDS mice of both genotypes and processed them for targeted NMR-based measures (Suppl. Fig. 4). Out of 25 metabolites, several were found to be significantly altered by CSDS in GR^lox/lox^ animals, including glutamate or aspartate (Fig. 4A, Suppl. File 4, Table S1-S3), which metabolites are used as biomarkers in human studies. These changes were not observed in CSDS-exposed GR^astroKO^ mice. In turn, the sole detected measure contrasting genotypes regarding the response to CSDS was a glutamate/glutamine ratio, an indicator of glutamate turnover operated by glutamine synthetase^31^. This data showed that CSDS induced metabolic effects in the brain, and revealed GR control of glutamate turnover in astrocytes.

**Fig. 4.**
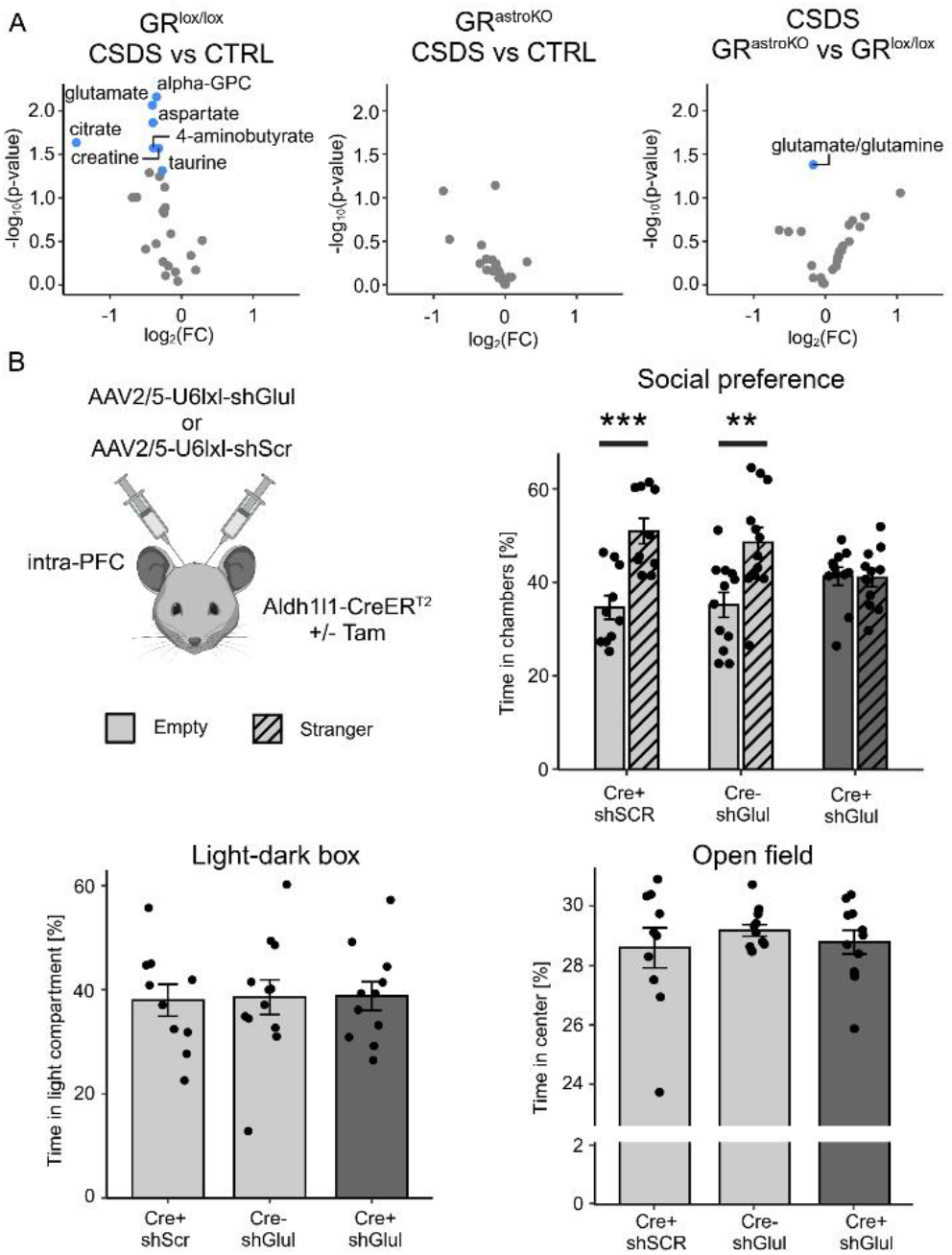
Astrocyte-specific metabolic pathways control circuit-specific behavioral traits. **A**. Volcano plots summarizing targeted NMR analysis of metabolites in cortical extracts from GR^astroKO^ and GR^lox/lox^ mice. Differential metabolites in pair-wise comparison are highlighted in red (upregulated) and blue (downregulated) **B**. Astrocyte-specific knock-down of Glul in the mouse PFC leads to altered social preference, without affecting anxiety. ** p<0.01,*** p<0.005, two-way ANOVA with post-hoc comparison.

Next we modeled the deficiency of the hallmark shared in human and mouse, i.e. glutamate turnover in the PFC and we examined its relevance for behavior. We performed a conditional knock down (KD) of *Glul* in the PFC astrocytes. A cassette containing shRNA targeting *Glul* or respective scrambled sequence (*Scr*) controlled by Cre-dependent U6 promoter (U6lxl) was delivered through AAV2/5 vector to the PFC of Aldh1l1-CreER^T2^ mice, followed by TAM injections. Mice with *Glul* KD (Cre+/sh*Glul*) and two control groups (Cre+/sh*Scr* and Cre-/sh*Glul*) were indistinguishable in light-dark box and open field tests, indicating unaffected exploration and anxiety phenotypes. In contrast, we detected significant differences in the time spent on social interactions, the behavior which relies on the intact function of the PFC^34,35^ (Fig. 4B, Suppl. File 4, Table S4-S6). This data shows that selective manipulation of the astrocyte-specific pathway controlling glutamate turnover is sufficient to elicit discrete traits of depressive-like behavioral spectrum, i.e. impaired social interactions.

### Intact GR signaling in astrocytes is required for behavioral effects of CSDS

Finally, we examined the role of GR signaling in astrocytes for CSDS-induced alterations of behavior. No effects of the GR elimination from astrocytes were observed in baseline behavioral measures, i.e. open field, light-dark box and 3-chamber social preference test (Suppl. Fig. S5). The exposure to CSDS of animals with intact GR resulted in significant decrease of the time spent in the light compartment and the time spent in the interaction chamber in respective tests. The elimination of the GR from astrocytes prevented these effects (Fig. 5A, B).

**Fig. 5.**
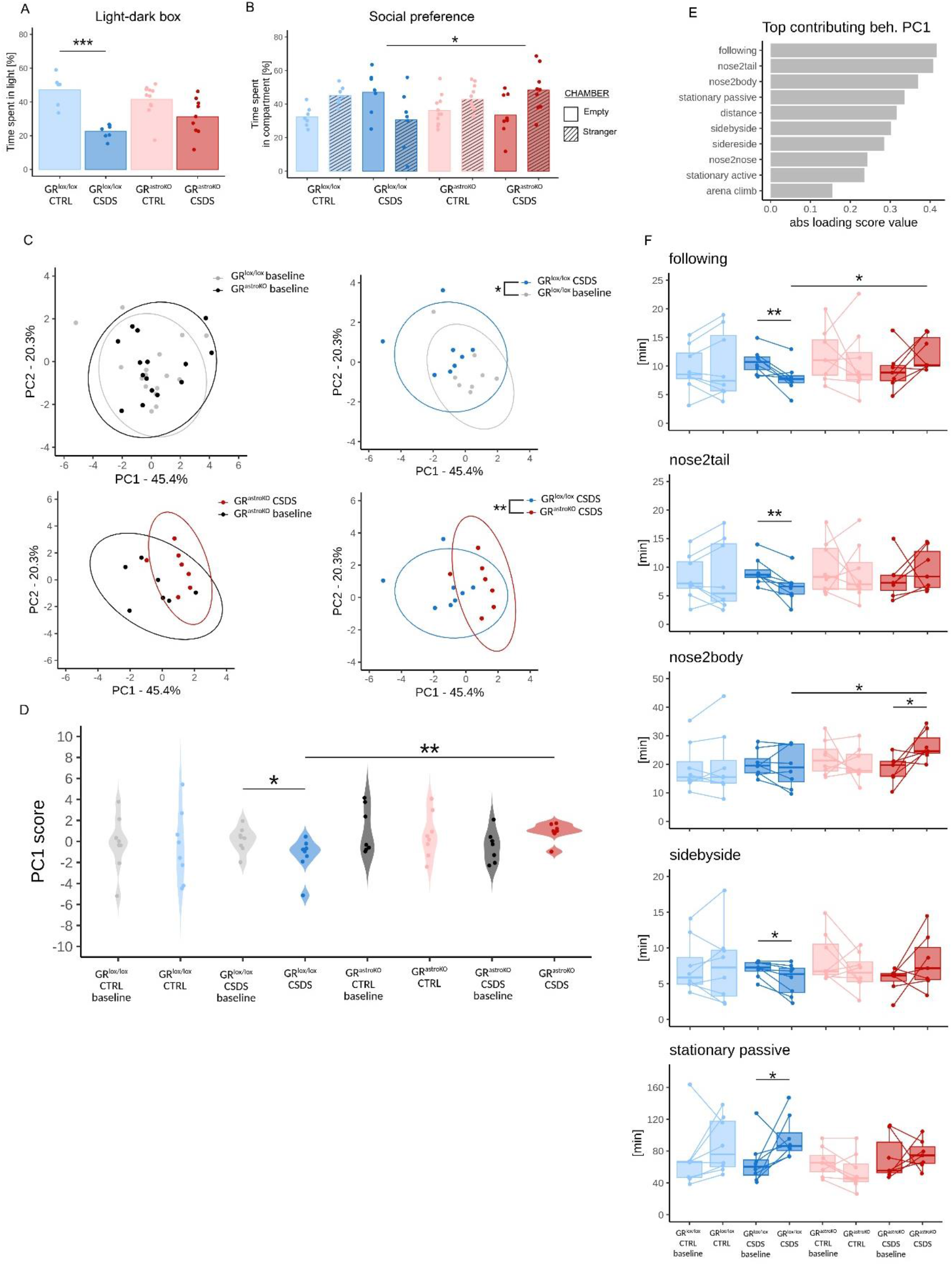
GR signaling in astrocytes mediates complex behavioral effects of CSDS. **A, B**. Graphs summarizing the results of classical behavioral tests performed after CSDS: **A**, light-dark box test and **B**, 3-chamber social preference test; p***<0.001; p*<0.05. **C-F**. Graphs summarizing the results of Social Box analysis performed before and after CSDS. **C**. PCA of behavioral traits before and after CSDS, assessed by deepOF supervised pipeline. Top-left graph shows cumulative results for GR^lox/lox^ and GR^astroKO^ animals at baseline. **D**. PC1 scores for groups in C; p*<0.05, p**<0.01. **E**. Top contributing behaviors influencing variation of the data, according to absolute loading scores of PC1. **F**. Top individual behavioral traits plotted before and after CSDS for each of 4 groups, assessed by deepOF supervised pipeline; p*<0.05.

After completing classical tests, we switched to an unbiased, automated analysis of the complex behavior. In this set of experiments, GR^lox/lox^ and GR^astroKO^ mice were placed in a square 50 cm x 50 cm arena and allowed to interact with a stranger conspecific. The analysis of animal postures was performed on video recordings (Sleap.ai), followed by machine learning - supported software (deepOF) which translated the data to multiple behavioral parameters (Suppl. Methods). This analysis did not reveal differences of baseline behavior between the GR^lox/lox^ and GR^astroKO^ males (Fig. 5C, grey vs black; Suppl. Fig. S5).

After baseline measures, animals from both groups were randomly assigned to CTRL (housed in pairs, light shades) or CSDS treatment (dark shades). Upon CSDS completion, all animals were exposed to the same conspecific as in baseline measures, and video recordings were analyzed as previously. The principal component analysis (PCA) of pre-vs post-treament data revealed significant effects of CSDS on complex behavior of GR^lox/lox^ mice, which were abolished in GR^astroKO^ mice (Fig. 5C, D). In line with this observation, the PCA of pre- and post-treatment data revealed that CSDS had differential impact on GR^lox/lox^ and GR^astroKO^ mice (Fig. 5D, dark blue vs dark red). Detailed analysis revealed that various parameters of social behavior contributed most to the observed variance (Fig. 5E). Statistical significance was observed in GR^lox/lox^ mice for the impact of CSDS on ‘nose2tail’, ‘following’, ‘sidebyside’ and ‘stationary passive’ parameters, which were abolished in GR^astroKO^ mice. In contrast, ‘nose2body’ was increased after CSDS in GR^astroKO^ mice, but not in GR^lox/lox^ mice (Fig. 5F). This data indicates complex role of astrocytic GR in mediating behavioral effects of CSDS, with some parameters being blunted, and some enhanced, suggestive for differential engagement of specific neural circuits.

Overall, the data from the rodent model indicated that GR signaling regulated transcriptional reprograming of astrocytes upon CSDS and these changes matched metabolic and behavioral outcome of the CSDS.

## DISCUSSION

This work reveals neurobiological mechanisms associated with depressive-like symptoms engaging defined signaling pathway, cell type, brain region, physiology and behavior. The BA25 plays a key role in negative emotion processing and was reported hypermetabolic in human depression^23,36^ and primate models of anhedonia^37^. Antidepressive therapies, such as BA25-targeted deep brain stimulation or ketamine, were shown to rescue BA25 metabolic hyperactivity, correlated with its antidepressant effects^37,38^. This region is engaged in coordinating stress response^29^ and highly sensitive to glucocorticoids^30^. Our study points to a crucial contribution of astrocytes to molecular and behavioral neuropathological phenotypes associated with severe mental disorders.

We show a transcriptional alterations of non-neuronal compartment as a key cellular deficit of BA25 in a subset of suicide completers, previously reported to display reduced expression of astrocyte-specific markers across the brain^18^. Our data extends previous findings by showing that 1) astrocytes’ reprogramming is a major cellular hallmark of BA25 in this cohort, 2) these changes engage the GR-dependent transcriptional network and 3) are accompanied by profound reprogramming of oligodendrocyte lineage. Through applying an astrocytic nuclei enrichment protocol, we were able to capture a detailed profile of BA25 astrocytes in a subpopulation of ‘low expressors’. This approach revealed that transcriptional networks in astrocytes crucially contribute to the aberrations of physiological processes which are hallmarks of depression: glutamate homeostasis^39^, protein trafficking^40^, lipid metabolism^41^, neural cells development^42^, and synaptic transmission^43^. The identification of GR as a main transcription factor regulating the gene network in ‘low expressors’ and mice exposed to chronic stress suggests that astrocytes may be a primary target of stress-induced, glucocorticoid-mediated effects in the brain.

Glucocorticoid resistance is among the most common systemic phenotypes in MDD^*44*^, which may explain physiological symptoms accompanying MDD, i.e. disruption of glucoregulatory mechanisms^*45*^ and sleep disturbances^*46*^. GR-controlled transcriptional networks, which may predict antidepressant response^11^, are highly heterogenous across organs^*47*^ and cell-specific within tissues. Our study deciphers that GR network regulates metabolic and synaptic genes in astrocytes. Interestingly, while the relationship between aberrant GR signaling and metabolism is well described in peripheral organs^*48*^, the brain is unique, as some of common metabolites, e.g., glutamate or glutamine, play specific roles in neuronal communication. Astrocytes play central role in glutamate homeostasis and energy metabolism in the brain. Neurobiological correlates of MDD include aberrant metabolism in several brain regions controlling affective behavior (e.g. PFC^*38*^), such as altered profile of glutamate metabolites^*49*^, an imbalance in excitatory/inhibitory neurotransmission^*50*^, aberrant glucose utilization^*51*^ and mitochondrial dysfunction^*52*^. Our data reveals downregulated expression of crucial molecular components of these pathways in astrocytes of ‘low expressors’, pointing to dysregulated GR network in astrocytes as a driver of neuropsychiatric phenotype. Importantly, normalization of metabolic parameters was shown to correlate with a positive outcome of antidepressant therapy^*53*^.

Glucocorticoids were also shown as powerful regulators of synaptic plasticity in physiology^54^ and under stress^55^. Previous studies showed that these effects may be due to cell-specific effects of GR in distinct population of neurons^*56*^ and glia^57^. In previous work, we showed that astrocytes are a prominent site of GCs action in the brain^57-59^. Here we show that astrocyte-specific GR ablation prevented transcriptional, metabolic and behavioral effects of chronic stress, supporting the hypothesis of astrocytes as the cellular locus relevant for central effects of glucocorticoids. Of note, an *in vivo* pharmacogenomic survey revealed that the regulation of GR-dependent genes enriched in astrocytes is a shared feature of psychoactive compounds, including antidepressants^60^. Together, these data highlight the importance of further research on antidepressants actions engaging astrocytic GR network.

Several studies reported decreased number of astrocytes in various brain regions of MDD cases. However, these studies exploited markers of reactive astrocytes. Considering the lack of evident signs of cell death, it cannot be ruled out that the decrease of specific proteins is rather due to molecular reprogramming of astrocytes which lose their functional identity. On the molecular level, we showed a dramatic downregulation of genes encoding astrocyte-specific glutamate transporter, GLT1 (SLC1A2), and a key enzyme mediating glutamate turnover, glutamine synthetase (GLUL) in BA25 of ‘low expressors’. These proteins were consistently reported as downregulated also in other brain areas in deceased cases with MDD diagnosis^17,18,20,61,62^. Causality experiments revealed that the pharmacological or genetic blockade of GLT1 in the BA25 led to increased activation of the BA25 and induction of anhedonic behavior in primates^37^; analogous data were obtained in rodent PFC^63-65^. In turn, pharmacological blockade of Glul in the PFC has led to inducing behavioral traits of depression in mouse^66^. Here, we provide a genetic evidence that the reduction of *Glul* in astrocytes elicited alterations of PFC-controlled behavior, i.e. abolished social preference, without detectable changes in tests measuring anxiety. Hence, primary dysfunction of glutamate metabolic pathway operated by astrocytes is sufficient to elicit circuit-specific phenotypes of depression.

In this work, we revealed dozens of pathways (GO BP terms) which expression differentiated human BA25 Cx43+ nuclei from CTRL and MDD, and overlapped with pathways differentiating PFC astrocytes from CTRL and CSDS-exposed mice. For many of these DEGs there already exists a rich body of evidence supporting their role at the synapse, such as regulation of synapse number (e.g. *Chrdl1*^*67*^, *Ptn*^*68*^, *Megf10*^*69*^) or synaptic transmission (e.g. *Slc1a2*^31^, *Glul*^31^, *Grin2c*^*70*^). In turn, many of identified genes encode key enzymes engaged in basic physiological processes, such as water transport (e.g. *Aqp4*^*71*^) or ion homeostasis (*Atp1b2*^*72*^), and crucial metabolic pathways, such as cholesterol synthesis (e.g. *Hmgcs1, Hmgcr*^*73*^) or energy metabolism (e.g. *Me1*^*74*^). The richness of genes altered in astrocytes provided in our study shall be exploited as a handle to manipulate astrocytes in order to restore their full operational capacity, a putative way of reversing synaptic and metabolic deficits in depression. This approach may be particularly useful in expanding a portfolio of biomarkers enabling patients stratification^75^ and development of more precise therapeutic strategies of psychiatric conditions^76^.

## AUTHORS CONTRIBUTIONS

Mi.S. and B.H. conceived of the project. S.D., B.Z., P.Z., D.D.P. V.D., T.K., B.H., and M.S. designed the study. M.S. wrote the manuscript with S.D., A.H., P.Z. and B.Z. contribution. S.D. established and performed the nuclei isolation protocol from human samples, refined with the help of C.V. and N.L., and performed initial analysis. G.T. selected and provided well-characterized human brain samples. S.D., N.L., M.K., V.B. and M.S. designed the RNA-sequencing experiments. D.H., M.P, and M.K. performed the RNA-sequencing data processing and S.D., A.H., M.P., D.H., Z.S., S.M.K., M.K. and M.S. performed the RNA-sequencing data analysis and interpretation. L.D. and F.G. performed the BrainTrawler analysis. S.D. established the protocol of isolating astrocytes from mouse samples and performed the isolation of mouse astrocytes with L.B. and C.M. assistance. S.D., P.Z., L.B., and C.M.P. performed the CSDS protocol, guided by H.S. and C.P instructions. V.D. and M.A. performed animal viral surgeries. P.H. and P.G. performed measures of tissue metabolites. L.B. performed the light-dark box and 3-chamber social interaction tests and analyzed the data with P.Z. contribution. B.Z. and P.Z. designed and installed the setup for automated analysis of mouse behavior and collected the data. B.Z. established the analytical workflow for automated analysis of behavior, to which M.V.S contributed by sharing his expertise and data. B.Z., P.Z., Z.S. and M.S. performed the analysis of complex behavior.

## ACKNOWLEDGMENTS

S.D., C.M.P., L.B., D.D.P., V.D., M.A., T.K., B.H. and M.S. are grateful to Dr. C. Tidona for his vision of a BioMed X Institute, to Ms. Y. Stappenbeck and Mr B. Reader for their support and members of BioMed X for creating an inspiring work environment. A.H., P.Z., B.Z, P.H. and M.S. are grateful for members of the AstroGroup for constructive discussions and feedback, and to J. Pierwoła and P. Krężel for help with IHC analysis. A.H., P.Z., B.Z., M.V.S. and M.S. acknowledge the support from the National Science Centre grant nr 2021/41/B/NZ3/04099 ‘AstroSyCo’ and HE Twinning ‘SAME-NeuroID: Standardized approaches to modelling and examination of neuropsychiatric disorders’. M.P., M.K. and M.S. acknowledge the support from the National Science Centre grant nr 2022/45/B/NZ5/03188 ‘GRtraits’. P.H., M.V.S. and M.S. acknowledge the support of collaborative grant from the Deutsche Forschungsgemeinschaft and the National Science Centre grant nr 2023/05/Y/NZ4/00124 ‘Astromics’.

