## Supplementary material for "Glucocorticoid receptors mediate reprogramming of astrocytes in depression": Material and methods

**HUMAN data.**

**Human samples.**

All procedures involving post-mortem samples of human brain were performed under the permission of the The Douglas Mental Health University Institute Research Ethics Board (Project number: IUSMD-17-43). Brain samples were dissected from BA25 by trained neuroanatomists at the Douglas Bell Canada Brain Bank and stored at -80˚C. Samples were originating from male subjects of Caucasians of French-Canadian descent and sudden death controls (N = 12), where the cause of death was determined by the Quebec Coroners' office. Cases (N = 15) met the criteria for MDD and died by suicide. Healthy controls did not have evidence of any axis I disorders. Care was taken to match the samples for the age, postmortem interval (PMI), pH, and refrigeration delay, whenever possible. Details on the subjects' information are listed in Suppl. File 1, Table 1.

**Fixation**

Hundred mg of frozen tissue was thawed on ice for 2 min. and was fixed with pre-cooled fixative solution (1% formaldehyde, 0.2 U µl−1 RNasin in PBS) for 8 min. on ice. Then, the fixative solution was discarded, and the tissue was washed with 1 ml of staining buffer (0.5% (wt/v) RNase-free BSA, 0.2 U µl − 1 RNasin in RNase-free PBS) by centrifugation at 400 × g for 5 min. at 4˚C.

**Homogenization**

The tissue was divided into two parts with a pre-chilled scalpel, and each half was transferred into a Dounce homogenizer containing a cold, freshly prepared homogenization buffer (250 mM sucrose, 25 mM KCl, 5 mM $MgCl_2$, 10 mM Tris, 1 µl DTT, 1x Protease inhibitor, 0.4 U/ µl RNAsin, 0.2 U/ µl SUPERase In, 01 % (v/v) Triton X-100 in nuclease-free water). Homogenization was initiated by applying 5 strokes using the loose pestle and completed with the tight pestle for another 10 strokes. The homogenates from two halves were combined and filtered using the cell strainer (40 µm pore size). Finally, the nuclei were pelleted by centrifugation at 1000 × g for 10 min. at 4˚C, and the supernatant was removed.

**Debris removal**

The pellet was resuspended with 250 µl of staining buffer and another 250 µl of 50% Optiprep solution (50% Optiprep with 42 mM sucrose, 25 mM KCl, 5 mM $MgCl_2$, 10 mM Tris pH 8.0 in nuclease-free water) was added on top and mixed gently to make 25% Optiprep-nuclei mixture. In a new tube, 500 µl of 29% Optiprep solution (29% Optiprep with 125 mM sucrose, 75 mM KCl, 15 mM $MgCl_2$, 30 mM Tris pH 8.0 in nuclease-free water) was placed, and the 25% Optiprep-nuclei mixture was layered on top. Next, two layers were obtained by centrifugation at 10,000 rpm for 30 min at 4˚C. Then, the supernatant was discarded without disturbing the pellet. Lastly, the pellet was resuspended with 1 ml of staining buffer and continued directly with immunostaining.

**Immunostaining**

Nuclei were blocked with staining buffer for 15 min. on ice, then incubated with 1000 µl of staining buffer containing primary antibodies for 1 h at 4˚C on a tube rotator. Nuclei were labelled with monoclonal anti-CX43 (2.5 µg/ml, Thermo-Fisher 13-8300), while the isotype control sample was labeled with mouse IgG1, kappa monoclonal antibody (2.5 µg/ml, ab91353). Nuclei were washed with staining buffer by centrifugation at 400 × g for 8 min. at 4˚C. Next, nuclei suspension was incubated with the secondary antibody coupled to Alexa 488 (4 µg/ml, Thermo-Fisher A-11029) for 45 min. at 4˚C on a tube rotator. Finally, after a washing step, the pellet was resuspended with 500 µl of staining buffer containing Hoechst (10 ng/ml), and samples were kept on ice until fluorescence-activated cytometry (FACS).

**Flow Cytometry**

Nuclei were analyzed and sorted using a FACS system (BD FACS Aria III) and associated software (BD FACSDiva 8.0.1) in the facility of the Zentrum für Molekulare Biologie der Universität Heidelberg (ZMBH). The particles' overall event rate was set up to 200 - 4,000 events/second, the sample loading chamber was kept at 4˚C, and a 100 µm nozzle was selected for sorting. First gating was performed using side and forward scatter channels for excluding debris and doublets. Intact nuclei were selected by sub-gating on Hoechst (405 nm excitation), constituting a fraction of 'Hoechst+ nuclei'. Subsequently, a subfraction of 'CX43+ nuclei' was defined based on the fluorescence intensity (488 nm excitation) compared to negative controls (isotype and secondary antibody controls). Next, size-based selection was performed, and 10-15% of smallest nuclei were excluded (Supp. Fig. S1). Selected nuclei populations were sorted into collection tubes containing 100 µl of PKD buffer (Qiagen) and stored at -80˚C until RNA extraction.

**RNA Extraction, Reverse Transcription, qPCR Analysis**

Total RNA was extracted using RNeasy FFPE Kit (Qiagen, 73504) with an additional elution step at the end; RNA was eluted twice with the same eluate (20 µl) to increase the total amount. Eluted RNA was collected into 1.5 ml non-stick, RNase-free microfuge tubes (ThermoFisher, AM12450). For quality control, 2 µl of

RNA was set aside in PCR strip tubes (Sigma, Z316156-1PAK) and, together with the rest of the eluted RNA, was stored at -80˚C. Quantity and quality assessments of RNA were done using 2100 Bioanalyzer Instrument (Agilent G2939BA), with RNA 6000 Pico Kit (Agilent, 5067-1513).

cDNA was synthesized with the SuperScript III system (ThermoFisher, 18080051) using oligo(dT) priming following the manufacturers' guidelines in the PCR Thermocycler T100 (Bio-rad, 1861096). qPCR experiments were run using the Applied Biosystems QuantStudio 3 (ThermoFisher).

qPCR analysis was accomplished with commercial TaqMan Probes or custom SYBR Green-based probes. Taqman probes specific for human species were synthesized and ordered following the suppliers' selection criteria (ThermoFisher). All the probes were carrying a FAM reporter. For experiments employing SYBR Green, custom primers were designed based on the following criteria: i. primers were spanning exon-exon junction, ii. the envisaged size of the amplicon was between 70-120 bp, and iii. primers had a low probability of self-dimerization, which were analyzed using the OligoAnalyzer Tool (IDT). The amplification efficiency of primers provided by the commercial supplier (Eurofins) was tested before the experiment, and the ones with 90-120% efficiency and a single peak on the melting curve were used for further experiments, employing the Fast SYBR Green Master Mix kit (Fisher Scientific, 4385612).

**RNA-seq data processing and differential expression analysis of experimental data – fig. 1A**

The quality of NGS data was verified using FastQC (v. 0.11.8). FastQC reports were aggregated with MultiQC (v1.10). The raw reads were aligned to GRCh38.p13 Ensembl (v. 102) using Hisat2 (v. 2.2.1). The gene abundances were calculated using htseq-count (v. 0.11.2) and GTF file from Ensembl (v. 102) as reference. Principal component analysis was performed with prcomp function from stats (v. 3.6.3) R package for various sets of genes. $Log_2$(CPM) density plots were created for various sets of data (raw and filtered data). Differential gene expression analysis between CTRL and MDD groups was done with glmQLFTest function from edgeR (v. 3.36.0) R package on filtered data (filterByExpr function with default parameters). The analysis was performed twice – separately for Hoechst and Cx43 samples and only using protein-coding genes. Hoechst sample genes were filtered for p-value < 0.05 and created a list of 843 differentially expressed genes (Suppl. File 1, Table S2). Cx43 sample genes were filtered for FDR < 0.1 and created a list of 67 differentially expressed genes – Cx43 list (Suppl. File 2, Table S1).

**Allen Brain Map & Braintrawler – fig. 1D**

Allen Brain Map Database consists of RNA sequencing data of single nuclei isolated from six areas of human cortex: middle temporal gyrus (MTG), anterior cingulate gyrus (CgGr), primary visual cortex (V1C), primary motor cortex (M1C), primary somatosensory cortex (S1C) and primary auditory cortex (A1C). The sequencing results were aligned to exons and introns in the GRCh38.p2 reference genome using the STAR algorithm, and aggregated intron and exon counts at the gene level were calculated. Gene expression matrix was downloaded from https://portal.brain-map.org/atlases-and-data/rnaseq/human-multiple-cortical-areas-smart-seq. Matrix contains one row for every cell in the dataset and one column for every gene sequenced. The values of the matrix represent total reads (introns + exons) for that gene (column) for that cell (row).

Specificity was computed with Braintrawler by dividing the mean gene expression in the cortex for a cell type by the mean expression in cortex over all cell types. In total the paper has 19 cell types (for 6 regions, so we see 19x6=116 cell type clusters in the paper) so the maximum cell type specificity is also 19. Abundance is computed with total reads count normalized by log(CPM(exons + introns)). Genes were filtered for specificity > 3 separately for each cell type creating 9 lists of genes specific for each cell type – Hodge lists.

**Cell-specific enrichment of DEGs in 'Hoechst+' fraction – fig. 1E**

Each of cell type specific lists was overlapped with Hoechst list by comparing shortnames (Suppl. File 1, Table S4-S12). Resulting number of overlapping celltype-specific, protein-coding genes from Hodge (with specificity > 3) with significant protein-coding genes from Hoechst (p-val < 0.05) (Z) alongside with number of overlapping protein-coding Hodge genes (all measured genes) with protein-coding Hoechst genes (all measured genes) (Grand Total - GT), number of Hodge celltype-specific genes (present in GT list, excluding list Z) (A), number of significant protein-coding genes in Hoechst list (present in GT list, excluding list Z) (B) were used to perform hypergeometric test (Suppl. File 1, Table S14). Bonferroni correction was applied to account for multiple comparison problem. All statistically significant sets and "unclassified" gene set were visualized by plotting $log_2$(FC) and mean $log_2$(CPM). Mean $log_2$(CPM) was computed as an average of $log_2$(CPM) of Hoechst control samples. "Unclassified" gene set consists of all the genes that were significant in Hoechst but not present in any cell type specific list (Suppl. File 1, Table S13).

**Over-representation analysis – fig. 1F**

ORA was done using Enrichr tool (https://maayanlab.cloud/Enrichr/) with overlaps of DEGs in 'Hoechst+' nuclei fraction and cell type specific lists. Results from ChEA 2022 – Transcription Factor Targets database were sorted by combined score. Combined score is the product of $\log_{10}$ of the p-value from the Fisher exact test and odds ratio. The odds ratio is computed using this formula:

oddsRatio = (1.0 * a * d) / max(1.0 * b * c, 1)

where: a are the overlapping genes, b are the genes in the annotated set - overlapping genes, c- are the genes in the input set - overlapping genes, and d- are the 20,000 genes (or total genes in the background) - genes in the annotated set - genes in the input set + overlapping genes. Top 10 sets were shown on the figures (Suppl. File 1, Table S15-S18).

**Rank-rank hypergeometric overlap – fig. 1C**

Rank-rank hypergeometric overlap was performed with RRHO (v. 1.34.0) R package. Genes on the X axis were ranked with $\log_{10}$(p-value) and genes on the Y axis were ranked with cell type specificity. Colors represent the exponents of hypergeometric p-values calculated for each point of the heatmap. Step size was 100 genes.

**DEGs and GR-dependent genes overlap – fig. 2D**

Venn diagram showing the overlap of differentially expressed genes in Cx43 samples (FDR<0.1) with GR-dependent genes from Carter et al. Mouse gene names from Carter were mapped to human gene names using file provided from Mouse Genome Informatics (https://www.informatics.jax.org/) (Suppl. File 2, Table S2)

**$\log_2$(FC) vs abundance plot – fig. 2C**

Significant genes were visualized by plotting $\log_2$(FC) and mean $\log_2$(CPM). Mean $\log_2$(CPM) was computed as an average of $\log_2$(CPM) of Cx43 control samples.

**Over representation analysis – fig. 2B**

Network was created with Metascape tool (https://metascape.org/) with significant protein-coding Cx43 genes as the input. Analysis was performed for Gene Ontology: Biological Process database. Results visualized with Cytoscape (v. 3.10.1) (Suppl. File 2, Table S3-S4).

**Gene Set Enrichment Analysis – fig. 2A**

GSEA was performed using clusterProfiler R package (v. 4.2.2.) for Gene Ontology – Biological Process database for Cx43 samples filtering for gene set size 5-200. Gene were ranked by formula:

$-\log_{10}$(p-value) × sign($\log_2$(FC)). Results were sorted by rank which is the position in the ranked list at which the maximum enrichment score occurred. Top 10 results were shown on the figure (Suppl. File 2, Table S5).

**MOUSE data.**

**Animals and housing conditions**

All animals experiments were conducted under the approval by the responsible Animal Care and Use Committees of either Regierungspräsidium Karlsruhe ethics committee (Karlsruhe, Germany) or Local Ethics Committee of the Institute of Immunology and Experimental Therapy PAS (Wrocław, Poland).

Behavioral tests were performed on males originating from breeding B6;FVB-Tg(Aldh1l1-cre/ERT2)1Khakh/J[1] transgenic line (Aldh1l1-CreER$^{T2}$) with he transgenic line carrying critical exons of gene encoding the glucocorticoid receptor (*Nr3c1*) flanked by loxP sites (Nr3c1tm2Gsc[2]). All animals were bred at C57Bl/6J background. Mice from cohort 1 were 2-3 months old, while mice from cohorts 2 and 3 were 6-8 months old at the beginning of experiments. Mice were group-housed in standard, individually ventilated IVC cages. Male CD-1 (Jackson Labs) were 8-10 months old ex-breeders, housed individually. As an unknown conspecifics, male mice with C57Bl/6J background were used (wild-types, WT): for 3 chamber social interaction test and for social boxes (3-6 months old). All mice were held under controlled temperature (22 ± 2°C) and humidity (± 45-65%) conditions with a 12/12h reversed light/dark cycle (lights off from 7 AM to 7 PM). During conducting the experiment all mice had ad libitum access to food and water. Experimental assay was planned to minimize the number of mice studied and to prevent any unnecessary stress to animals.

**Viral vector production and AAV injections**

Primers for amplifying the shRNA against *Glul* or its scrambled version were cloned to HpaI/XhoI site of the pSico plasmid (Addgene, #11578) carrying Cre-dependent version of the U6 polymerase, followed by CMV-EGFP (FwdshGlul: TGCATGTCACTAAAGCGGGCTCAAGAGGCCCGCTTTAGTGACATGCTTTTTC; RevshGlul: TCGAGAAAAAGCATGTCACTAAAGCGGGCCTCTTGAGCCCGCTTTAGTGACATGCT; FwdshScr: TGCAGGCACGTGGCAATCATTCAAGAGATGATTGCCACGTGCCTGCTTTTTC; RevshScr: TCGAGAAAAAGCAGGCACGTGGCAATCATCTCTTGAATGATTGCCACGTGCCTGCA). Afterwards, XbaI/KpnI sites were used for cloning the cassette to a helper plasmid. Vectors of the AAV2/5 serotype were produced by a commercial vendor (Vector Biolabs) with a reported titer of 2.5 x $10^{12}$. Experiments were carried out using 10-12-week-old males of Aldh1l1-CreER$^{T2}$ transgenic line, bred at the animal facility of the University of Heidelberg. All mice were housed under standard light cycle conditions (12:12h, lights off 19:00-07:00) with ad libitum access to food and water. To inject AAV-shGlul or AAVshScr, mice were anaesthetized with a sleeping mix (Medetomodin, 60 µl of 1 mg/ml solution; Midazolam, 160 µl of 5 mg/ml solution; and Fentanyl, 40 µl of 0.05 mg/ml solution; volume µl = 3 x [mouse weight] in g), received analgesia (Carprofen, s.c., 5 mg/kg) and eye protection (Bepanthen), and placed in a stereotactic frame. After skin disinfection, a cut was made at midline, connective tissue removed, and injections sites marked (AP: 2mm, ML: ± 0.2mm from Bregma). Injection needle was lowered at DV: -3 mm and after 30 s to allow the settling of the tissue, 50 nl of the vector solution was administered at 1 nl/s, followed by 3 min. rest. Subsequently, the procedure was repeated at DV: -2.5 mm, - 2 mm, and -1.5 mm). The procedure was repeated in the second hemisphere. The skin was sutured with non-absorbable vicryl threat. After injection completion, mice received a waking mix (Atipamezol, 30 µl of 5 mg/ml solution; Flumazenil, 30 µl of 0.1 mg/ml solution, Naloxon, 180 µl of 0.4 mg/ml; and 360 µl of saline; double volume of the sleeping mix). Mice were then placed on a heating pad until full recovery.

**Tamoxifen and dexamethasone (DEX) administration**

To excise the loxP/loxP site by Cre recombinase, mice were injected intraperitoneal (IP) with 2 mg of TAM (Sigma Aldrich) for 5 consecutive days. TAM was dissolved in 1:9 ratio in ethanol (Chempur) and sunflower seed oil (Sigma Aldrich) at a final concentration of 10 mg/mL.

**Chronic Social Defeat Stress paradigm (CSDS)**

The CSDS paradigm was conducted according to the published protocol[3] with a few modifications. Custom-designed cages with transparent, perforated dividers were used (Tecniplast, Buguggiate, Italy). CD-1 male mice were screened for max. 3 consecutive days for probability to attack an unknown C57Bl/6J male mice and the most aggressive subjects („aggressors") were selected for the CSDS procedure. To prevent bite wounds of tested mice, lower incisors of CD-1 mice were trimmed (Erbrich, Tuttlingen) under brief isoflurane anaesthesia every second day. On the day prior to the first social defeat session, an experimental mouse was introduced to one of the two compartments in the aggressor's homecage to allow sensory contact, without physical interactions between animals. After 24 hours, experimental mouse was placed in the chamber with CD-1 and allowed for 10 minutes of direct contact or at least 60 sec of attack. Subsequently, tested mouse was checked for wounds and re-introduced to the home cage of unknown aggressor. Social defeat paradigm lasted for 15 consecutive days, on which body weight and health status of each animal were monitored daily. Control groups were housed in pairs, in cages with transparent, perforated dividers, allowing only sensory contact.

**Classical behavioral tests – Fig. 4B, 5A,B**

The battery of behavioral tests was conducted a minimum of 4-weeks after the first tamoxifen injection. Mice were allocated to the CTRL and CSDS groups, ensuring even distribution of animals regarding initial body weight (BW) and anxiety score (based on the Open Field Test performed pre-CSDS). All mice underwent at least 1 hour of adaptation to the room and test environment before each behavioral experiment. Each test was performed in the same time points/conditions for both experimental groups (7 AM - 11 PM; ZT1 - ZT5). Tests were held under video-based rejestration and analyzed with Ethovision XT15 Software (RRID:SCR_000441, Noldus Information Technology, Wageningen, Netherlands).

**Open field test**

Open field test was performed at baseline, in an empty arena (41.5 cm x 41.5 cm x 50 cm), build out of transparent material under constant lighting conditions of approximately 60 lux. Mice were allowed to explore arena freely for 5 minutes, in which time spent in the center zone (sec) and distance travelled (cm) were measured (Suppl. File 4, Table S5).

**Light-dark box test**

To assess anxiety-related behavior in mice, the light-dark box test was performed. Behavioral arena was made out of opaque material and consisted of a light (56 cm x 25 cm x 25 cm, 500 lux) and dark compartment (12 cm x 25 cm x 25 cm, < 5 lux), connected with center-located opening (3.5 cm x 3.5 cm). Mice were introduced to the dark chamber of arena and were able to explore it freely for 10 minutes. For each mouse time spent in both light and dark compartments was measured and analysed (Suppl. File 2, Table S4).

**Three chamber social interaction test**

Three chamber social interaction test was conducted as previously reported[4], under red light conditions. The behavioral arena was made out of transparent plexi material and consisted of three, equally sized compartments (45 cm x 20 cm x 25 cm), connected with eachother with openings (3.5 cm x 3.5 cm) located in the center of separators. In each outer compartment a wire-meshed cage was placed. The paradigm included two-step approach: a habituation phase, in which experimental mice was introduced to the middle compartment and allowed to explore freely arena with empty wire-meshed cages for 5 minutes. Then, mouse was gently guided and locked in the center chamber, while a novel conspecific („stranger", C57Bl/6J background male mouse) was placed randomly in one of the wire-meshed cages. The second testing phase started when experimental mouse was allowed to explore freely all three chambers and lasted 10 minutes. To quantify preference for social approach behaviors the time spent in each compartment and interaction zone (around the wire-meshed cages) was measured (Suppl. File 2, Table S6).

**Social Box**

Spontaneous individual and social behavior was studied in a custom-designed arenas (boxes), to represent semi-naturalistic environment. The setup incorporated 4 square, open arenas 50 x 50 x 60 cm (LxWxH), with installed feeders and holders for water-bottles. Bedding and enrichment (cotton rolls) were added, in accordance with 3R guidelines. Arenas were separated with each other with thick, black curtains, allowing for sound and light restriction, coming from possible outside sources. During the experiment arenas were illuminated with infra-red light of 940nm wavelength in order to conduct studies under complete darkness. We have used standard and cheap 48 diodes-IR-plates (3W DC12V 300mA) available through global stores. Two IR plates were used per one box and the distance between the plates and floor of the arena was 85cm. Animals in social boxes were recorded with Basler daA1920-160um cameras paired with Fujinon 6mm f1.8 C125-0618-5M lenses. Distance between the lens and the floor surface was 85cm to ensure proper capture of the whole arena and to minimize possible barrel distortion of the lens. Launching and recording the experiment was carried out with Bonsai.rx software, which allowed for simultaneous registering of the image in all boxes at the same time, for the same exact amount of time. Experiments were carried out in pairs. One animal was of $GR^{lox/lox}$ or $GR^{astroKO}$ genotype, while the second animal was an unknown conspecific (C57Bl/6J background). Each WT animal had tail markings to ensure visual discrimination between subjects, necessary for the subsequent workflow. Social box evaluation was conducted at first 4 weeks after TAM injections, serving as a baseline for post-CSDS session performed after 14 days of CSDS. We recorded all animals for 4 hours, at the beginning of the active phase. Videos were saved in .avi format, with 648 x 600 dimensions (WxH) recorder at 25 frames per second, in greyscale.

**Behavior analysis - Fig. 5C-F**

Once acquired, all videos were processed with SLEAP.ai[5] software for pose-estimation and subsequently with deepOF[6] software for actual behavioral prediction. Because deepOF requires at least 11 annotation points on a single animal, we established annotations in accordance with guidelines at deepof.readthedocs.io. After pilot analyses, we decided to place additional 2 annotations on a tail of each animal to boost the performance of machine learning models, as.more annotation points allow for better association between them and render better inference on supplied videos after training. The analyses was performed on the personal computer with NVIDIA Tesla A100 40GB on board. To achieve valid pose-estimation, we have created a multi-animal top-down model using 148 labelled frames from 8 separate videos and unet as a backbone.

Parameters for Centroid Model Configuration were as follows:

Data section: Validation fraction – 0.1; Input Scaling – 0.75; Crop Size – Auto.

Optimazation section: Batch Size – 4; Epochs – 100; Initial Learning Rate – 0.0001; Stop Training on Plateau – disabled; Online Mining – disabled.

Augmantation section: Rotation selected, with Rotation Min Angle as -180 and Rotation Max Angle as 180; Scaling disabled; Uniform Noise disabled; Gaussian Noise disabled; Contrast enabled with Contrast Min Gamma of 0.5 and Contrast Max Gamma as 2; Brightness enabled with Brightness Min Val of 0 and Brightness Max Val of 10.

Model section: Backbone – unet; Stem Stride selected as None; Max Stride – 16; Filters – 16; Filters Rate – 2; Middle Block selected; Up Interpolate selected;

Heads section: centroid; Anchor Part – Center; Sigma – 2.5; Output Stride – 2.

Parameters for Centered Instance Model Configuration were as follows:

Data section: Validation fraction – 0.1; Input Scaling – 1; Crop Size – 160.

Optimization section: Batch Size – 4; Epochs – 200; Initial Learning Rate – 0.0001; Stop Training on Plateau – disabled; Online Mining – enabled, with Min Hard Keypoints of 1 and Max Hard Keypoints selected as None.

Augmentation section: Rotation selected, with Rotation Min Angle as -180 and Rotation Max Angle as 180; Scaling disabled; Uniform Noise disabled; Gaussian Noise disabled; Contrast enabled with Contrast Min Gamma of 0.5 and Contrast Max Gamma as 2; Brightness enabled with Brightness Min Val of 0 and Brightness Max Val of 10.

Model section: Backbone – unet; Stem Stride selected as None; Max Stride – 16; Filters – 24; Filters Rate – 2; Middle Block selected; Up Interpolate selected;

Heads section: centered_instance; Anchor Part – Center; Sigma – 2.5; Output Stride – 4.

After creating the model, prediction of all annotation points was carried out with following Inference Pipeline (Multi-Animal-Top-Down Pipeline):

Max Instances – 2 (each for individual animal in the box); Tracker (cross-frame identity) Method - simple; Max number of tracks – 2 (each for individual animal in the box); Similarity method – iou; Matching Method – Hungarian; Elapsed Frame Window – 5; Robust quantile of similarity scores selected as Use max (non-robust); Kalman filter-based tracking was deselected, as in this case it did not improve stability of animals ids; Nodes use for tracking – all 13 used; Connect Single Track Breaks was enabled.

Although highly successful, SLEAP.ai would occasionally switch identities of the animals, particularly during inference (e.g. during fights between subjects and in cases where animals were very close to each other for a longer period of time). Thus, proofreading and manual correction of identities was necessary to maintain valid input for behavioral assessment with deepOF. Subsequently, all prepared input files were provided for deepOF workflow (please visit deepof.readthedocs.io for detailed guidelines). We have used supervised pipeline within deepOF and used 11 annotation points on the animal body to predict sets of individual and dyadic behavior parameters. Finally, the .csv files (delivering predicted behavior parameters) provided by deepOF, were processed with custom scripts written in Rstudio for final analysis and visualization.

**Statistics**

Statistical analysis was done and plots were created using Rstudio, ggplot2 package, svglite package and BioRender software. For classical tests, normality and homogeneity of variance was estimated with Shapiro−Wilk and Levene tests, followed by a two-way ANOVA and post-hoc tests. Tests were performed using GraphPad Prism version 9.4.0 for Windows, GraphPad Software, Boston, Massachusetts USA). For Social Box analyses, based on previous data[6] and our pilot analyses, data from first 10 minutes of video recordings were used. Data were tested for normality using Shapiro-Willk test with shapiro.test() function. For comparison of separate groups, the unpaired t-test was used. For comparison of behavioral alterations in the same group (baseline vs after CSDS), a paired t-test was used. Both tests were done with t.test() function. PCA analysis was performed with prcomp() function. Significance was considered at values: $p^{*}<0.05$; $p^{**}<0.01$; $p^{***}<0.001$.

**Immunofluorescence – Fig. 3B**

Immunohistochemistry staining was performed as previously described[7]. Mice were perfused intracardially with PBS, followed by 4% paraformaldehyde (PFA) buffered with phosphate buffered saline (PBS) and brains were postfixed in 4% PFA and stored in PBS. Coronal sections from $GR^{lox/lox}$ and $GR^{astroKO}$ mice obtained with vibratome (70-µm-thick). Free-floating sections were rinsed in PBS (3 times), incubated with 10% normal goat serum in PBST (0.2% Triton X−100 in PBS) for 90 min at room temperature (RT). Sections were then incubated at RT with antibodies: polyclonal anti-S100β (1:250, 287 004, Synaptic Systems) or a monoclonal anti-NeuN (1:200, MAB377, Merck Millipore) and a monoclonal anti-GR (1:200, 12041S, Cell Signalling) diluted in PBS-Tx containing 1% normal serum. After 48h, sections were washed in PBS and incubated for 90 min at RT with fluorophore-conjugated secondary antibodies (A11073, A11036, A11029, Thermo- Fisher), diluted 1:1000 in PBS-Tx containing 1% normal serum. Afterwards, sections were washed in PBS, incubated for 10 min at RT with Hoechst (1:1000 dilution in PBS, 62249, ThermoScientific) and embedded in VECTASHIELD® Antifade Mounting Medium (H-1000, Vector Laboratories). Imaging was performed with a fluorescent confocal microscope (Zeiss Cell Observer SD).

**Astrocyte isolation.**

For RNAseq analyses, mice were sacrificed at the beginning of the active phase (ZT1-ZT2), brains were extracted and hippocampi and prefrontal cortices were dissected. Astrocytes were isolated according to the published protocol[8], with modifications. Freshly dissected mouse tissues were homogenized using the Adult Mouse Brain Dissociation kit (Miltenyi) with implemented modifications. Brain tissues were minced with a pre-cooled scalpel on ice (omitted for PFC), transferred into the tube containing pre-heated Enzyme mix 1 (50 µl Enzyme P + 1900 µl Buffer Z), and incubated for 10 min. at 37˚C. Next, Enzyme mix 2 (10 µl Enzyme A + 20 µl Buffer Y) was added, and the first round of trituration was performed (5 strokes, for whole-brain and cortex: 10 strokes) with an unpolished Pasteur pipette. After incubating for 5 min. at 37˚C, the second trituration was conducted (5 strokes, for whole-brain and cortex: 10 strokes) with a medium-sized fire-polished Pasteur pipette. Then, 1 ml of cell suspension was placed into a new tube and kept on ice. The remaining homogenate was incubated for 10 min. at 37˚C. Next, 125 U/ml of DNase I was added, and final trituration was done (5 strokes, for whole-brain and cortex: 10 strokes) using a small-sized fire-polished Pasteur pipette. Finally, the homogenate was filtered (the remaining 1 ml cell suspension kept on ice was applied as well), and the cells were pelleted by centrifugation at 300×g for 10 min. at 4˚C.

For PFC, pellets from the previous step were resuspended in 100 µl of PB buffer (DPBS, pH 7.2, and 0.5% BSA) and kept on ice. For hippocampi, debris removal was conducted according to the manufacturers' instructions with slight modification; the pellet from the homogenization step was resuspended in cold DPBS and mixed with a cold debris removal solution. Next, cold DBPS was gently overlayed to obtain clear two phases and centrifuged at 3000×g for 10 min. at 4˚C with full acceleration and full brake. The top two phases were discarded, cold DPBS was added to a final volume of 15 ml and was gently mixed by inverting the tubes 3 times. Cells were pelleted by centrifugation at 1000×g for 10 min. at 4˚C with full acceleration and full brake. Next, pellets were resuspended in PB buffer, and myelin removal beads (Myelin Removal Beads II kit, Miltenyi) were added, mixed by pipetting up and down, and incubated for 15 min. at 4˚C. Cells were washed with PB buffer and centrifuged at 300×g for 10 min. at 4˚C. Next, myelinated cells were removed through a magnetic separator using MS columns. Myelin-positive cells were retained in the column, and the flowthrough (myelin-negative cells) were collected and centrifuged at 300×g for 10 min. at 4˚C.

To isolate astrocytes, anti-ACSA-2 MicroBeads kit (Miltenyi) was used, precisely following the manufacturers' protocol. Positive selection was performed where astrocytes were magnetically labelled with the beads, retained within the column during the magnetic field, and eluted. Next, cells were pelleted by centrifugation at 300×g for 10 min. at 4˚C, washed with 500 µl of cold DPBS, and centrifuged again at 300×g for 5 min. at 4˚C. Finally, the pellets were resuspended in 350 µl of RLT buffer containing β-mercaptoethanol (10 µl of β-ME per 1 ml of RLT buffer) and stored at -80˚C.

**RNA extraction and library preparation**

RNA was extracted with RNeasy Plus Micro Kit (Qiagen). Eluted RNA (20 µl) was collected into 1.5 ml non-stick RNase-free microfuge tubes (ThermoFisher, AM12450), and stored at -80˚C. cDNA was prepared using a modified version of Smart-seq2 protocol. Nextera XT DNA Library Preparation Kit (Illumina) was used following the internal protocol of EMBL Genomics Facility. Maximal amount of RNA (2.4 µl) was employed to generate cDNA, and 0.2 ng of amplified cDNA was used with 20-40 ng/µl custom-made Tn5 enzyme (EMBL Genomics Facility) for the tagmentation of cDNA. Next, PCR amplification was performed for adapter-ligand fragments adding a unique pair of i5 and i7 adapters (Illumina) in the reaction. Lastly, RNA and library quality controls were done using Bioanalyzer system with RNA 6000 Pico Kit and Agilent High Sensitivity DNA kit. Samples were pooled (mix of 13-16 libraries in one tube) and sequenced using NextSeq 500 system (Illumina) at EMBL Genomics Facility (dual index, paired-end 2×75 bp run).

**RNA-seq data processing and differential expression analysis of experimental data – fig. 3A**

The quality of NGS data was verified using FastQC (v. 0.11.7). FastQC reports were aggregated with MultiQC (v1.10). The raw reads were aligned to GRCm38.p6 Ensembl (v. 99) using Hisat2 (v. 2.1.0). The gene abundances were calculated using featureCounts and GTF file from Ensembl (v. 99) as reference. Principal component analysis was performed with pca function from PCAtools (v. 2.4.0) R package. Differential gene expression analysis between 3 pairs: $GR^{lox/lox}$/CSDS vs $GR^{lox/lox}$/CTRL, $GR^{astroKO}$/CSDS vs $GR^{astroKO}$/CTRL, $GR^{astroKO}$/CSDS vs $GR^{lox/lox}$/CSDS was done with glmQLFTest function from edgeR (v. 3.36.0) R package on filtered data (with function filterByExpr from edgeR R package) for protein-coding genes for both tissue regions separately (hippocampus, prefrontal cortex). Genes were filtered for p-value < 0.05 and created a list of 413 differentially expressed genes for $GR^{lox/lox}$/CSDS vs $GR^{lox/lox}$/CTRL in Hipp, 240 for $GR^{astroKO}$/CSDS vs $GR^{astroKO}$/CTRL in Hipp, 941 for $GR^{astroKO}$/CSDS vs $GR^{lox/lox}$/CSDS in Hipp, 488 $GR^{lox/lox}$/CSDS vs $GR^{lox/lox}$/CTRL in PFC, 64 for $GR^{astroKO}$/CSDS vs $GR^{astroKO}$/CTRL in PFC and 345 for $GR^{astroKO}$/CSDS vs $GR^{lox/lox}$/CSDS in PFC (Suppl. File 3, Table S1-S6).

**Astrocyte-specific GR enrichment fig. 3C-E**

For each of the 3 comparisons: $GR^{lox/lox}$/CSDS vs $GR^{lox/lox}$/CTRL, $GR^{astroKO}$/CSDS vs $GR^{astroKO}$/CTRL, $GR^{astroKO}$/CSDS vs $GR^{lox/lox}$/CSDS overlap between significant genes in PFC, significant genes in Hipp and GR-dependent genes from Carter et al. was checked. Hypergeometric test was calculated for the overlaps between Hipp and Carter, PFC and Carter and Hipp and PFC. Background genes were selected as an overlap between all genes measured our experiment and genes measured in Carter et al. Bonferroni correction was applied to account for multiple comparison problem (Suppl. File 3, Table S7).

**Over representation analysis – GO: BP database - Metascape analysis – fig. 3F**

Network was created with Metascape tool (https://metascape.org/) with protein-coding, significant in $GR^{astroKO}$/CSDS vs $GR^{lox/lox}$/CSDS in PFC comparison genes as the input. Analysis was performed for Gene Ontology: Biological Process database (Suppl. File 3, Table S8-S9). Results visualized with Cytoscape (v. 3.10.1). Circles in bold represent GO terms that were also found significant in human MDD vs CTRL in Cx43 comparison (Suppl. File 3, Table S10).

**Heatmaps of differentially expressed genes belonging to selected GO BP Metascape clusters – fig. 3G**

For selected Metascape clusters Z-scores of genes belonging to each cluster of GO terms were shown on a heatmap. Heatmaps were constructed using heatmap3 function from heatmap3 R package (v. 1.1.9). Black and white bar shows the abundance of the genes.

**Tissue processing and NMR measures – Fig. 4A**

Mice were sacrificed at the end of the active phase (ZT23-ZT24) by cervical dislocation, and the whole brain was dissected into distinct regions, including the cortex, hippocampus, and cerebellum. All brain tissues were immediately frozen in liquid nitrogen and stored at -80°C until further analysis. The brain tissue was weighed into an Eppendorf tube, and 500 µL of LC-MS grade methanol (-80°C; Supelco, 1.06035) was added. Tissue lysis was performed using a Qiagen Retsch TissueLyser II at 30 Hz for 2 minutes, repeated twice. Between the two runs, samples were briefly cooled in liquid nitrogen. Subsequently, 250 µL of ultrapure water (Supelco, 1.01262) was added, and the samples were vortexed for 5 minutes, followed by brief centrifugation.Next, 600 µL of the sample was transferred to a separate Eppendorf tube, mixed with 400 µL of chloroform (Sigma-Aldrich, 650741), vortexed for 5 minutes, and centrifuged at 18,000 RCF for 15 minutes at 4°C. The upper polar layer (400 µL) was collected into a 5 mL Falcon tube, and 250 µL of ultrapure water was added to adjust the eutectic point of the mixture. Finally, samples were freeze-dried overnight. For the remaining protein interphase and nonpolar chloroform layer, 800 µL of methanol was added, and the sample was centrifuged at 18,000 RCF for 15 minutes. The supernatant was discarded, and the protein pellet was washed three times briefly with water before being solubilized in 0.1 M NaOH for normalization using the Pierce™ BCA Protein Assay.

NMR samples and raw spectra were prepared according to standard Chenomx protocols. The lyophilized extracts were dissolved in 200 µl of phosphate buffer and data acquired were on a Bruker 700 MHz instrument equipped with a cryoprobe and Topspin 3.6 software. The data were processed and analyzed using Chenomx versions 10, 11 and 12. Spectra were referenced to dss, baseline corrected and the pH adjusted within the Chenomx software.

Lyophilized experimental samples were reconstituted in 200 µL of the prepared Chenomx I NMR buffer (1.2063 g of $HK_2PO_4$, 413.2 mg of $H_2KPO_4$, 12.26 mg of $NaN_3$, 1.074 mg of DSS-$d_6$, and 3.072 mg of imidazole were dissolved in $D_2O$ to a final volume of 100 mL in a volumetric flask. This resulted in a 200 mM phosphate buffer at pH 7.3, containing 0.03% $NaN_3$, 0.5 mM DSS-$d_6$, and 20 mM imidazole buffer). The samples were mixed at 500 rpm for 1 minute, followed by centrifugation at 1,000 g for 1 minute to recover the supernatant. A volume of 160 µL of the resulting solution was transferred to a clean 3 mm NMR tube for further analysis.

NMR spectra were acquired using the Chenomx protocol for 1D $^1H$ NMR, as detailed in their reference documentation (https://www.chenomx.com/wp-content/uploads/2018/10/N007.pdf). A Bruker Avance III 700 MHz NMR spectrometer equipped with a TCI cryoprobe and Topspin 3.6 software was used to acquire the NMR data. Data were collected using the METNOESY pulse sequence with the following parameters: Temperature 298 K, acquisition time (AQ) 4 s, mixing time (D8) 0.1 s, initial delay (DE) 0.01 s, pre-saturation delay (D1) 0.99 s, steady state scans (DS) 16, spectral width (SW) 12 ppm. After tuning, matching and shimming of the first sample, the frequency of the residual water resonance was determined and used for all experiments in the batch (O1). The 90 degree $^1H$ pulse (P1) was also determined for the first sample and assumed to be a constant for each batch of experiments carried out in the same buffer. The number of transients (scans, NS) was optimized for the first sample (ns = 64, 256 and 1024 tested). The number of transients was set to 1024 in order to monitor as many metabolites as possible. The increased number of scans improved the number of observable metabolites from less than 20 to about 34 although several were at the edge of detection and not all peaks were observed.

The acquired spectra were analyzed using Chenomx NMR Suite for quantitative metabolite profiling. The Bruker fids were loaded and processed using Chenomx Processor. The automatically phased spectra were manually adjusted, a line broadening of 0.5 Hz was applied and the baseline corrected using the Whittaker spline method and manually picked baseline points. The spectrum was calibrated to 0 ppm for the dss peak and the pH was calibrated on the imidazole peaks. The corrected spectra were transferred to Chenomx Profiler. Here, peak assignments were made on the basis of the software suggestions and concentration guidelines. The peak positions were manually adjusted for a better fit and concentration fitting was made using the linear least squares fitting routine within the software. The optimized fits were manually checked and adjustments made where significant NMR peak density was unaccounted for, with another round of least squares fitting. The compound table was exported to an excel workbook for further statistical analyses.

1 Srinivasan, R. *et al.* New Transgenic Mouse Lines for Selectively Targeting Astrocytes and Studying Calcium Signals in Astrocyte Processes In Situ and In Vivo. *Neuron* **92**, 1181-1195 (2016). https://doi.org:10.1016/j.neuron.2016.11.030
2 Tronche, F. *et al.* Disruption of the glucocorticoid receptor gene in the nervous system results in reduced anxiety. *Nat Genet* **23**, 99-103 (1999). https://doi.org:10.1038/12703
3 Sigrist, H., Hogg, D. E., Senn, A. & Pryce, C. R. Mouse Model of Chronic Social Stress-Induced Excessive Pavlovian Aversion Learning-Memory. *Curr Protoc* **4**, e1008 (2024). https://doi.org:10.1002/cpz1.1008
4 Naert, A., Callaerts-Vegh, Z. & D'Hooge, R. Nocturnal hyperactivity, increased social novelty preference and delayed extinction of fear responses in post-weaning socially isolated mice. *Brain Res Bull* **85**, 354-362 (2011). https://doi.org:10.1016/j.brainresbull.2011.03.027
5 Pereira, T. D. *et al.* SLEAP: A deep learning system for multi-animal pose tracking. *Nat Methods* **19**, 486-495 (2022). https://doi.org:10.1038/s41592-022-01426-1
6 Bordes, J. *et al.* Automatically annotated motion tracking identifies a distinct social behavioral profile following chronic social defeat stress. *Nature communications* **14**, 4319 (2023). https://doi.org:10.1038/s41467-023-40040-3
7 Tertil, M. *et al.* Glucocorticoid receptor signaling in astrocytes is required for aversive memory formation. *Translational psychiatry* **8**, 255 (2018). https://doi.org:10.1038/s41398-018-0300-x
8 Batiuk, M. Y. *et al.* An immunoaffinity-based method for isolating ultrapure adult astrocytes based on ATP1B2 targeting by the ACSA-2 antibody. *J Biol Chem* **292**, 8874-8891 (2017). https://doi.org:10.1074/jbc.M116.765313
